# A novel role for the extraembryonic area opaca in positioning the primitive streak of the early chick embryo

**DOI:** 10.1101/2021.10.26.465879

**Authors:** Hyung Chul Lee, Claudio D. Stern

## Abstract

Classical studies have established that the marginal zone, a ring of extraembryonic epiblast immediately surrounding the embryonic epiblast (area pellucida) of the chick embryo is important in setting embryonic polarity by positioning the primitive streak, the site of gastrulation. The more external extraembryonic region (area opaca) was only thought to have nutritive and support functions. Using experimental embryology approaches, this study reveals three separable functions for this outer region: first, juxtaposition of the area opaca directly onto the area pellucida induces a new marginal zone from the latter; this induced domain is entirely posterior in character. Second, ablation and grafting experiments using an isolated anterior half of the blastoderm and pieces of area opaca suggest that the area opaca can influence the polarity of the adjacent marginal zone. Finally, we show that the loss of the ability of such isolated anterior half-embryos to regulate (re-establish polarity spontaneously) at the early primitive streak stage can be rescued by replacing the area opaca by one from a younger stage. These results uncover new roles of chick extraembryonic tissues in early development.

**Summary statement:** Two adjacent extraembryonic tissues, the area opaca and the marginal zone, interact to influence the polarity of the early chick embryo.

## Introduction

Before the start of gastrulation (formation of the primitive streak), the chick embryo is disc-shaped and comprises three concentric regions: an inner (central) disc called area pellucida (which includes all cells destined to contribute to the embryo proper), an outermost ring called the area opaca and an intermediate, narrow ring called marginal zone (Lee et al., 2020). A considerable body of classical work has implicated the latter, and especially its posterior portion (“posterior marginal zone”) in determining the site of primitive streak formation within the adjacent area pellucida (Bachvarova et al., 1998, Khaner and Eyal-Giladi, 1986, Khaner and Eyal-Giladi, 1989, Khaner et al., 1985, Shah et al., 1997, Skromne and Stern, 2001, Spratt and Haas, 1960, Torlopp et al., 2014, Bertocchini and Stern, 2012, Azar and Eyal-Giladi, 1979, Eyal-Giladi and Khaner, 1989). The posterior marginal zone is a strong signalling centre that expresses cVG1 (currently labelled GDF3 in the chicken genome), whose misexpression in the anterior marginal zone is sufficient to initiate formation of an ectopic primitive streak (Seleiro et al., 1996, Shah et al., 1997, Skromne and Stern, 2001). The rest of the marginal zone displays a posteriorly-decreasing gradient of expression of BMP4 and its targets such as GATA2, which can act as inhibitors of primitive streak formation (Bertocchini and Stern, 2012, Sheng and Stern, 1999). In the posterior area pellucida, two early targets of cVG1 signalling from the marginal zone are essential for initiating primitive streak formation at this site: cVG1 itself (Skromne and Stern, 2002) and another TGFβ superfamily member, NODAL (Bertocchini and Stern, 2002), which may act together as a heterodimer to induce mesendodermal fate (Montague and Schier, 2017, Opazo et al., 2019).

In contrast, the more peripheral region, area opaca, is generally believed to play a less active role in regulating embryonic patterning, its major functions at this early stage being providing a source of nutrition and maintaining the centrifugal tension of more central regions through adhesion of its edge to the vitelline membrane (Bellairs et al., 1967, Downie, 1976, New, 1959). Indeed, the embryo can generate a primitive streak even when the area opaca is removed, which had been interpreted to imply that the area opaca has no role on embryonic polarity (Spratt and Haas, 1960, Khaner et al., 1985).

The present study uncovers three separable roles of the area opaca in the regulation of polarity of the early embryo. First, it is able to induce a marginal zone (without polarity) when placed directly adjacent to the area pellucida. Second, it is able to induce posterior character on adjacent marginal zone and/or area pellucida embryo fragments. Finally, it appears to be responsible for the loss of the ability of isolated embryo fragments to form a primitive streak after the time of appearance of the endogenous primitive streak (stage HH2) (Bertocchini et al., 2004, Spratt and Haas, 1960).

## Results

### Molecular differences between anterior and posterior area opaca

Previous studies have revealed a few genes whose expression differs in anterior and posterior parts of the extraembryonic area opaca of the chick embryo. Among these, BMP4 and GATA2 are more strongly expressed anteriorly (Bertocchini et al., 2004, Sheng and Stern, 1999, Streit et al., 1998, Torlopp et al., 2014), whereas cWnt8C is expressed more strongly posteriorly (Hume and Dodd, 1993, Skromne and Stern, 2001). A recent study explored regional differences in different parts of the embryo more systematically using RNAseq (Lee et al., 2020). To uncover the most significant molecular differences between the anterior and posterior ends of the area opaca, we ranked the relative expression levels of these two regions. Table 1 lists several genes showing the highest level of differential expression – these observations suggest that the area opaca is polarized along the anterior-posterior axis.

**Table 1.** List of genes with polarized expression pattern in the area opaca (AO)

| **aAO > pAO** |  |  |  | **aAO < pAO** |  |  |  |
| --- | --- | --- | --- | --- | --- | --- | --- |
| **gene_name** | **aAO** | **pAO** | **fold change** | **gene_name** | **aAO** | **pAO** | **fold change** |
| TFAP2A | 111.733 | 69.121 | 1.616484 | PITX2 | 10.861 | 60.436 | 5.56479 |
| DHRS3 | 88.221 | 56.270 | 1.567812 | TBX6 | 10.448 | 52.940 | 5.06696 |
| GATA2 | 145.400 | 93.504 | 1.555007 | ASTL | 40.426 | 122.057 | 3.019285 |
| CYP26B1 | 62.195 | 43.950 | 1.41514 | LYPLA1 | 32.217 | 90.928 | 2.822361 |
| SMAGP | 94.706 | 67.391 | 1.405326 | LITAF | 54.256 | 127.402 | 2.348177 |
| ALS2CL | 53.905 | 39.205 | 1.374951 | GNOT2 | 66.512 | 150.297 | 2.259704 |
| ETS2 | 54.225 | 40.634 | 1.334466 | KPNA7 | 28.228 | 53.240 | 1.886098 |
| CMIP | 238.602 | 180.113 | 1.324735 | FZD7 | 33.256 | 61.600 | 1.852293 |
| BASP1 | 55.481 | 42.017 | 1.320461 | APLNR | 33.741 | 60.845 | 1.803274 |
| FAM53A | 99.004 | 75.178 | 1.316932 | FIBIN | 65.248 | 117.624 | 1.802714 |

a, anterior; p, posterior

### The area opaca can induce functional properties of the marginal zone

A considerable body of work spanning several decades has revealed a crucial role for the marginal zone (a belt of cells intervening between the embryonic, central area pellucida and the extraembryonic, outer, area opaca) in regulating the polarity of the early blastoderm by positioning the site at which the primitive streak will form (Khaner and Eyal-Giladi, 1986, Khaner and Eyal-Giladi, 1989, Khaner et al., 1985, Spratt and Haas, 1960, Bachvarova et al., 1998). The observations in the previous section that the area opaca is polarized raises the possibility that it may also influence the polarity of either the marginal zone, or the area pellucida (directly or indirectly).

First, we explored whether the area opaca can induce marginal zone properties from area pellucida cells and if so, whether the induced marginal zone is functionally polarized. To investigate this we ablated the marginal zone. This frees a larger piece of area opaca than is necessary to surround the area pellucida (in the absence of the intervening marginal zone), so we grafted area opaca strips lacking the posterior or the anterior parts (about 60° arc). We assessed the results by examining the expression of the marginal zone marker *ASTL* and the posterior marginal zone marker *cVG1* after 8 h (Fig. 1). The isolated area pellucida did not express *ASTL* (0/4; Fig. 1 E), confirming complete ablation of the marginal zone and suggesting that this region does not spontaneously regenerate. In contrast, a graft of area opaca (without the posterior region) induced a thin ring of *ASTL* expression in the adjacent area pellucida in some embryos (5/13 with expression) (Fig. 1 F). The induced expression of *ASTL* resembles that in the marginal zone of normal embryos (Fig. 1 D). In the normal embryo, strong expression of *cVG1* is initially (stage X-XI) concentrated in the posterior marginal zone, later followed by expression in the neighbouring posterior area pellucida (Fig. 1 A, G) and later in the primitive streak itself. When the area pellucida was cultured alone, weak expression of *cVG1* in the posterior region was maintained (Fig. 1 B and H). In contrast, a graft of area opaca (lacking the posterior part) induced expression of *cVG1* in the area pellucida either as a ring or at multiple sites (8/9 with ring-like or multiple sites of expression, 1/9 in the posterior region only) (Fig. 1 C, I).

**Figure 1.**
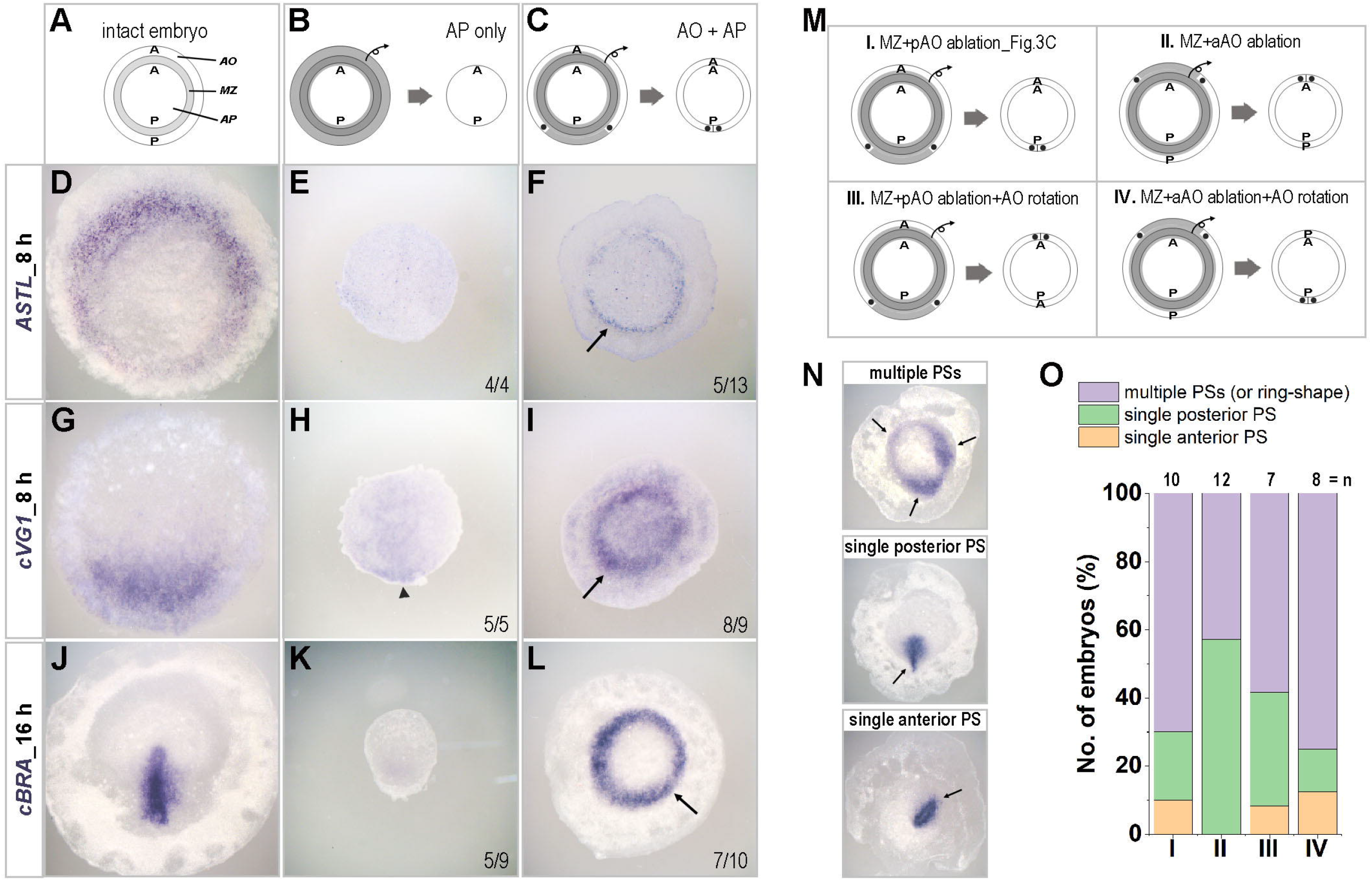
Induction of a marginal zone by the area opaca. (A, D, G, J) Expression of the marginal zone marker *ASTL* (D), of the posterior marginal zone marker *cVG1* (G) after 8 h culture, and of the primitive streak marker *cBRA* (J) after overnight culture, in normal embryos. AO, area opaca. MZ, marginal zone. AP, area pellucida. (B, E, H, K) Expression of the same markers in embryos after ablation of the area opaca and the marginal zone. Some weak expression of *cVG1* is observed in the posterior area pellucida (arrowhead, H), while neither *ASTL* nor *cBRA* is expressed (or very low expression, in case of *cBRA*). (C, F, I, L) Expression of the markers in embryos after ablation of the marginal zone and grafting area opaca. In many cases, all makers (including the posterior markers *cVG1* and *cBRA*) show multiple or ring-shape expression patterns (arrows). The proportion of embryos showing the morphology illustrated is indicated in each panel. (M-O) Effects of varying the orientation of the grafted regions or ablation of the area opaca. (M) Experimental design. A, anterior. P, posterior. (N) The resulting embryos could be assigned to three morphological classes. Arrows indicate primitive streak formation with *cBRA* expression. (O) Graph showing the incidence of each type of result.

To test whether expression of *cVG1* is followed by primitive streak and mesendoderm formation, we cultured the operated embryos overnight (16 h) and examined the expression of *cBRA*. Unlike normal embryos, where *cBRA* is expressed in the streak (Fig. 1 J), the area pellucida alone showed no (or faint) expression (5/9 with no expression, 4/9 with weak posterior expression) (Fig. 1 K). When the area pellucida was surrounded by a strip of area opaca lacking the posterior part (Fig. 1C), most embryos (7/10) showed multiple sites or ring expression of *cBRA*, 2/7 had a single posterior streak and 1/7 had a single streak arising anteriorly (Fig. 1 L). When the area pellucida was surrounded by a strip of area opaca lacking the anterior part (also a 60° arc), some embryos (3/7) also showed multiple foci or ring expression of *cBRA*, but a greater proportion (4/7) had a single posterior site of *cBRA* expression (Fig. 1 M-O, conditions I and II).

As a further test of the ability of the area opaca to influence polarity of the area pellucida, we combined these juxtaposition experiments with 180° rotation of the area opaca relative to the area opaca (Fig. 1 M-O, conditions III and IV). In both cases (rotation of area opaca lacking either its anterior or its posterior portion) the majority of grafts generated a ring or multiple sites of *cBRA* expression (7/12 and 6/8 respectively), and the remainder generated a single primitive streak positioned either posteriorly or anteriorly in the area pellucida (Fig. 1 M-O, conditions III and IV).

Together, these results show that the area opaca can induce a functional marginal zone in the area pellucida. However, the ring-shape, rather than polarized, expression of *cVG1* and *cBRA* in the majority of embryos receiving an area opaca graft suggests that induction of the marginal zone by the area opaca can be separated from establishment of polarity in the marginal zone.

### The area opaca can bias the polarity of an isolated anterior half-embryo

In the above experiments, the area pellucida itself has already experienced the influence of more peripheral tissues (marginal zone and perhaps area opaca) by the time the experiment starts. As a more unbiased assay, we turned to the anterior half of the blastoderm cultured by itself. As previously described, such a fragment will initiate primitive streak formation from either the left or the right posterior edge of the area pellucida, with equal frequency (Bertocchini et al., 2004, Spratt and Haas, 1960, Torlopp et al., 2014). Therefore this provides a “sensitized” system to assess subtle influences of the area opaca on polarity.

First, we ablated a small piece of posterior lateral area opaca from one side of the anterior half, whose polarity was then assessed by expression of *cVG1* (after 6 h) or *cBRA* (after overnight culture). Expression of *cVG1* was observed, now more frequently in the marginal zone of the opposite side to the ablation (Fig. 2 A, B, E). The marginal zone slightly anterior to the excision site (but not that immediately adjacent) showed weak expression of *cVG1* (Fig. 2 B). After overnight culture, *cBRA* expression was also biased to the opposite side to the excision (Fig. 2 A, C, D, E). These results suggest that the area opaca influences the polarity of an isolated anterior half-blastoderm.

**Figure 2.**
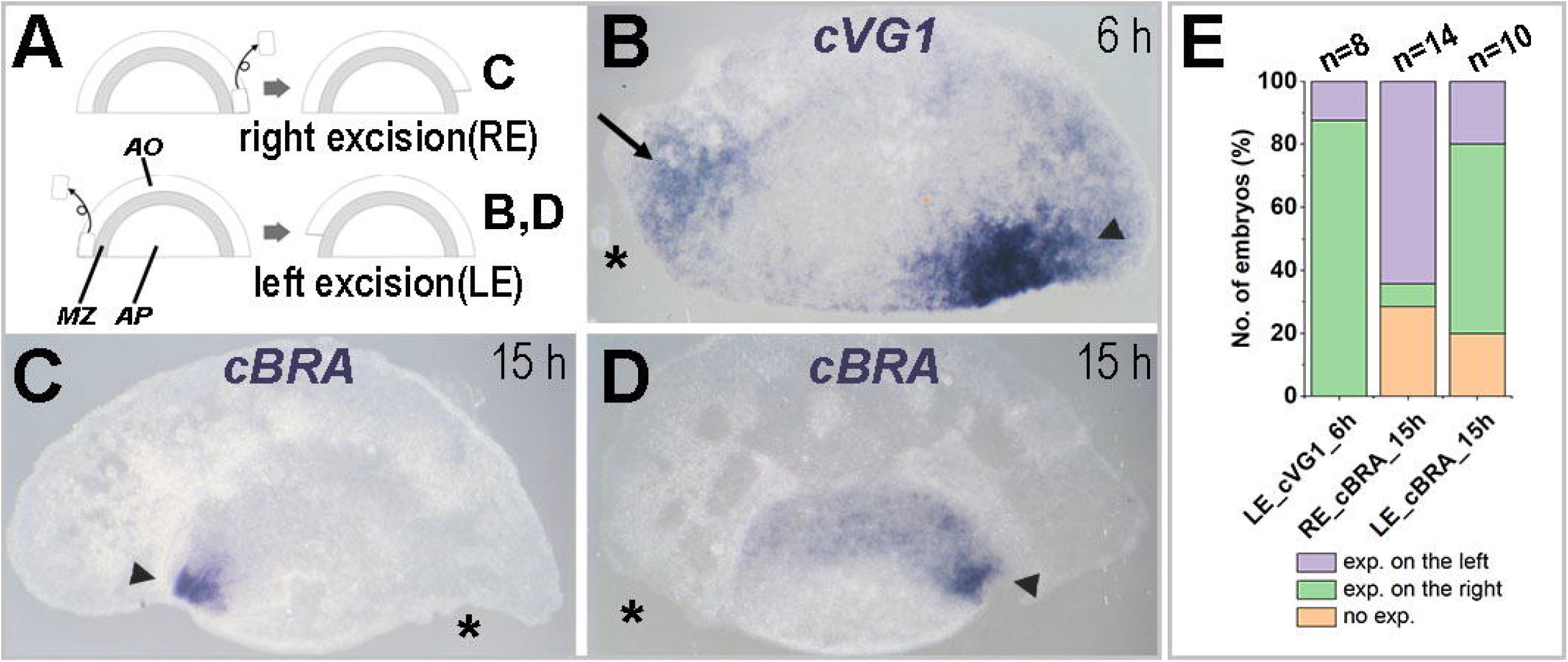
Excision of area opaca from an isolated anterior half-embryo biases *cVG1* expression and primitive streak formation. (A) Experimental design. Isolated anterior half-embryos are cultured after excision of either the right (RE; for C) or left (LE; for B, D) edge of the area opaca. AO, area opaca. MZ, marginal zone. AP, area pellucida. Embryos were processed for expression of *cVG1* after 6 h (B) or *cBRA* (primitive streak) after overnight culture (C, D). (B) After 6 h, *cVG1* is expressed more strongly at the contralateral side to the excision (arrowhead), but weak *cVG1* expression is also observed in the marginal zone lying immediately anterior to the excision site of area opaca (arrow). (C, D) After overnight culture, a single primitive streak forms from the contralateral edge of the area pellucida (opposite the excision site; arrowhead). (E) Summary graph showing the number of embryos with different expression patterns. Asterisks, site of excision.

### Primitive streak-inducing ability of the area opaca

To test whether the polarity of the area opaca itself exerts an influence on the site of primitive streak formation of the isolated anterior half, a piece of anterior area opaca was replaced by its posterior counterpart (from the other half of the embryo) (Fig. 3 A). The donor posterior half-embryo was examined for *cVG1* expression immediately after excision to ensure that the graft did not include the *cVG1* expressing region of the marginal zone (Fig. 3 B) (Shah et al., 1997). After short incubation (6 h), *cVG1* expression was observed in the marginal zone near the grafted donor tissue [7/10 (70%) with *cVG1* expression, 3/10 (30%) with no expression] (Fig. 3 C). After overnight incubation *cBRA* expression was observed near the grafted donor tissue in 7/19 embryos (37%), while the remaining 12 embryos (63%) had formed a streak from the left or right posterior edge, not near the donor tissue (Fig. 3 F). Control grafts (excision and replacement of the anterior area opaca from the same embryo) resembled simple isolated anterior halves in that the majority (6/7; 86%) had one site of *cBRA* expression either on the left or the right posterior edge; the remaining embryo had *cBRA* expression anteriorly, near the site of the graft (Fig. 3 E). To check for any contribution of the donor cells to the primitive streak, GFP-transgenic embryos (McGrew et al., 2008) were used as donors of posterior area opaca. Immunostaining with anti-GFP antibody revealed no contribution of donor cells to the marginal zone or the primitive streak after overnight incubation (3/8, Fig. 3 G; 5/8 had formed a primitive streak from the posterior edge but not near the graft). These results are consistent with the idea that the posterior area opaca emits inducing signals that can polarize the embryo and position the site of primitive streak formation, even in the presence of the endogenous marginal zone.

**Figure 3.**
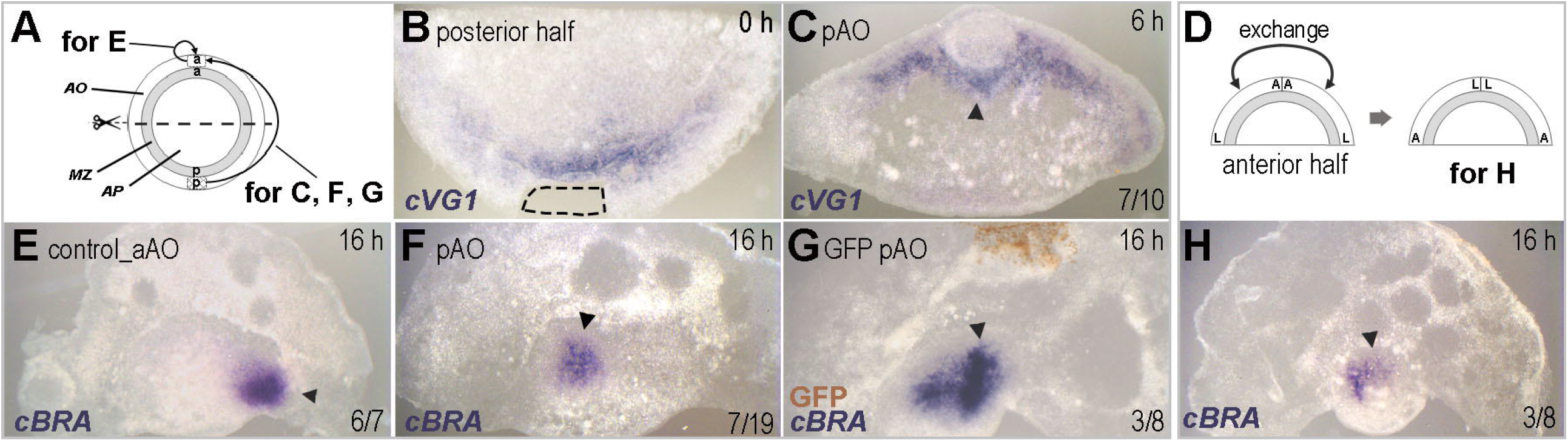
A graft of posterior area opaca onto an isolated anterior half-embryo induces *cVG1* expression and primitive streak formation. (A) Experimental design for B, C, E, F and G. AO, area opaca. MZ, marginal zone. AP, area pellucida. (B) The posterior piece of area opaca (not expressing *cVG1*) is used as the donor for grafting (dotted line). (C) After 6 h, *cVG1* is induced in the marginal zone adjacent to the graft (arrowhead). (E-F) After overnight culture, an ectopic primitive streak (*cBRA* expression) is generated near the graft (arrowhead, F), while control grafts (anterior area opaca) have no effect – the primitive streak forms from one edge of the isolated anterior half-embryo as it does in the absence of a graft (arrowhead) (E). (G) Using donor tissue taken from GFP-transgenic embryos reveals no cellular contribution of the graft to the induced ectopic primitive streak (arrowhead) (G). (D-H) When the area opaca of the isolated anterior half-embryo is cut in half and the left and right fragments exchanged (to swap the anterior and lateral aspects of the area opaca) (D), an ectopic primitive streak with *cBRA* is induced (arrowhead) (H). The proportion of embryos showing each illustrated morphology is indicated in each panel.

### The area opaca is responsible for the loss of the regulative ability of the primitive streak stage embryo

The regulative ability of an isolated anterior half-blastoderm is lost as soon as the primitive streak starts to form (Spratt and Haas, 1960, Streit et al., 1998) (Fig. 4 A, C, D, F, I), raising the question of whether the area opaca may be at least partly responsible. To test this, we grafted the anterior area opaca of a pre-primitive-streak stage embryo (stages EGK X-XI; “early”) to the inner region (area pellucida and marginal zone) of the anterior half of a primitive-streak stage host (stage HH 2-3; “late”), or vice versa (Fig. 4 C, G). When early area opaca was placed adjacent to a late inner anterior half, most cases showed *cBRA* expression (20/22), indicating that the early stage area opaca can rescue the regulative ability of a later stage anterior half (Fig. 4 C, F, I). In the converse experiment, when late anterior area opaca was placed next to a younger inner anterior half-blastoderm (area pellucida and marginal zone), the frequency of primitive streak formation was 62 % (13/21) (Fig. 4 G, J, I), which represents a reduction in frequency relative to anterior halves from early stage embryos with their own area opaca, which generate a primitive streak in 100% of cases (24/24) (Fig. 4 A, D, I). This decrease in frequency raises the possibility that the late area opaca exerts an inhibitory influence on primitive streak formation. To test this, we cultured the anterior half of early embryos in the absence of the area opaca (Fig. 4 H). They exhibited a similar reduction in frequency of primitive streak formation (Fig. 4 K, I), suggesting that the late area opaca does not gain inhibitory properties but rather loses its inducing, or polarity-promoting, functions. Together, our results reveal novel roles for the extraembryonic area opaca, including both the ability to induce a marginal zone, and a polarising influence that can result in positioning the site of primitive streak formation.

**Figure 4.**
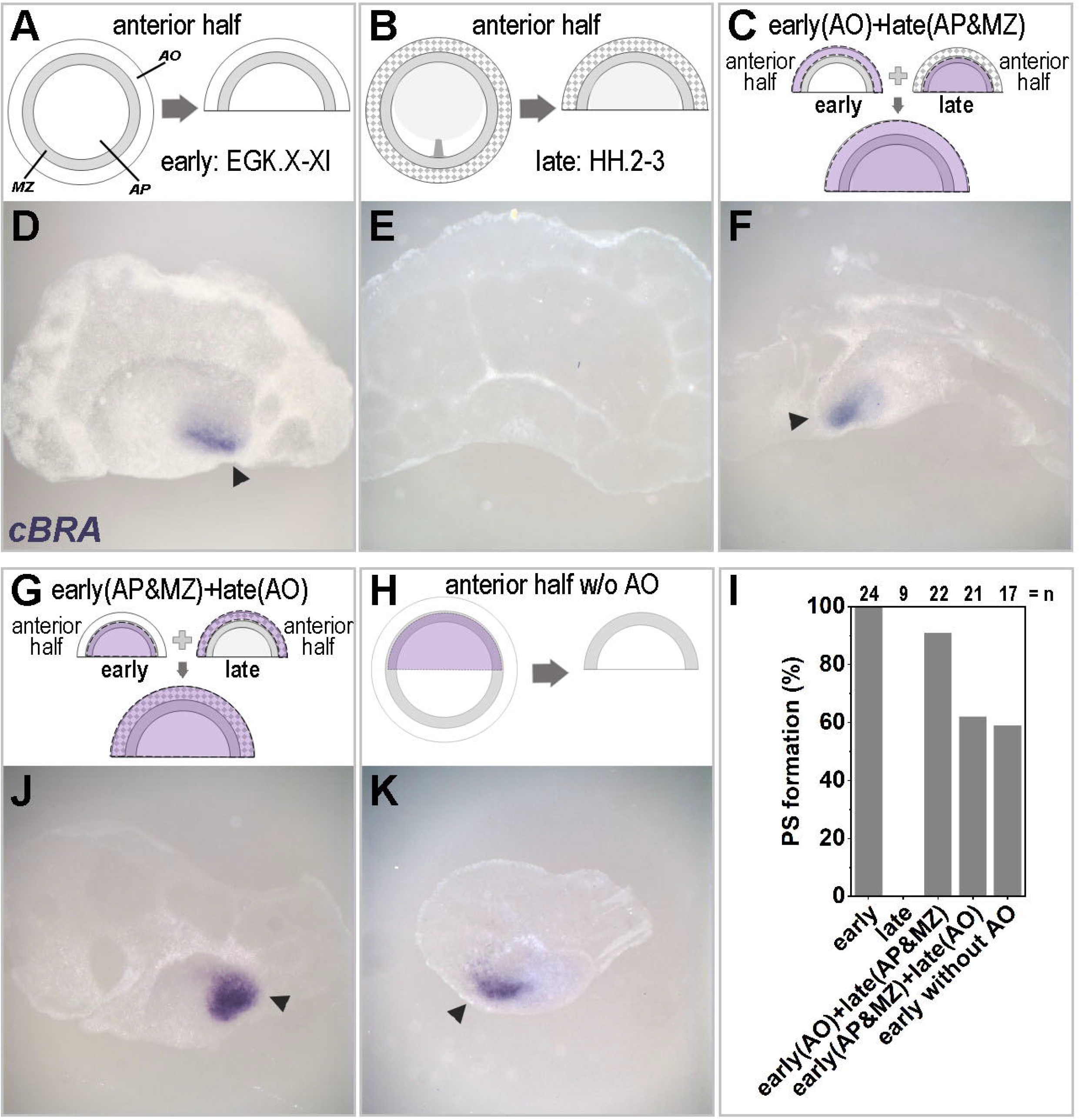
The early area opaca can rescue the loss of regulative ability of primitive streak stage anterior half-embryos. (A, D) The anterior half of pre-primitive-streak-stage (“early”) embryos spontaneously generates a primitive streak expressing *cBRA* (Arrowhead, E) from either the left or right posterior edge. AO, area opaca. MZ, marginal zone. AP, area pellucida. (B, E) In contrast, the anterior half of primitive-streak-stage (“late”) embryos can no longer generate a primitive streak. (C, F) Replacing the anterior area opaca of a late anterior half-embryo with the equivalent region from an early donor rescues the regulative ability of the late-stage embryo fragment, generating a primitive streak (arrowhead). (G, J) Conversely, grafting late anterior area opaca onto a younger host reduces the regulative ability of the latter to the same level as anterior half-embryos deprived of the area opaca at the same stage (H, K, I). (I) graph showing quantification of the results.

## Discussion

The early chick embryo (prior to formation of the primitive streak) could be viewed as consisting largely of a single layer of epithelial cells that is continuous across its three concentric regions: the area pellucida at the centre (containing all prospective embryonic cells), the area opaca at the periphery, and an intermediate thin ring of cells, the marginal zone lying between the previous two (Duval, 1889, Eyal-Giladi and Kochav, 1976, Pander, 1817, Pasteels, 1940, Stern, 2004, Vakaet, 1970, Kochav et al., 1980) (Fig. 5A). Beneath this single ectodermal layer are different tissues that largely do not contribute to the embryo: the single-cell-thick hypoblast and endoblast underlying the area pellucida (the former does carry germ cell precursors), and a thick, multi-layered spongy arrangement of large yolky cells, called the germ wall, underlying the area opaca and marginal zone (Stern, 1990, Stern and Downs, 2012, Vakaet, 1970, Stern, 2004). Marking the boundary between area pellucida and marginal zone at the posterior edge of the former is a sharp ridge of cells, Koller’s sickle (Bachvarova et al., 1998, Callebaut and Van Nueten, 1994, Izpisúa-Belmonte et al., 1993, Koller, 1882), which protrudes ventrally beneath the ectoderm and provides a bridge between this and the underlying hypoblast/endoblast. Although the marginal zone boundaries are not obvious morphologically in intact embryos other than at this posterior domain, brushing off the edges of the germ wall towards the area opaca reveals a continuous ring where the germ wall is not attached to the overlying ectoderm, corresponding to the region that had been called marginal zone in the classical literature (Eyal-Giladi and Kochav, 1976, Kochav et al., 1980). A recent study (Lee et al., 2020) uncovered the first molecular marker restricted to this entire region: ASTL, encoding an astacin-like metalloendopeptidase, confirming the existence of a distinct anatomical region that surrounds the entire area pellucida as proposed by previous observations.

**Figure 5.**
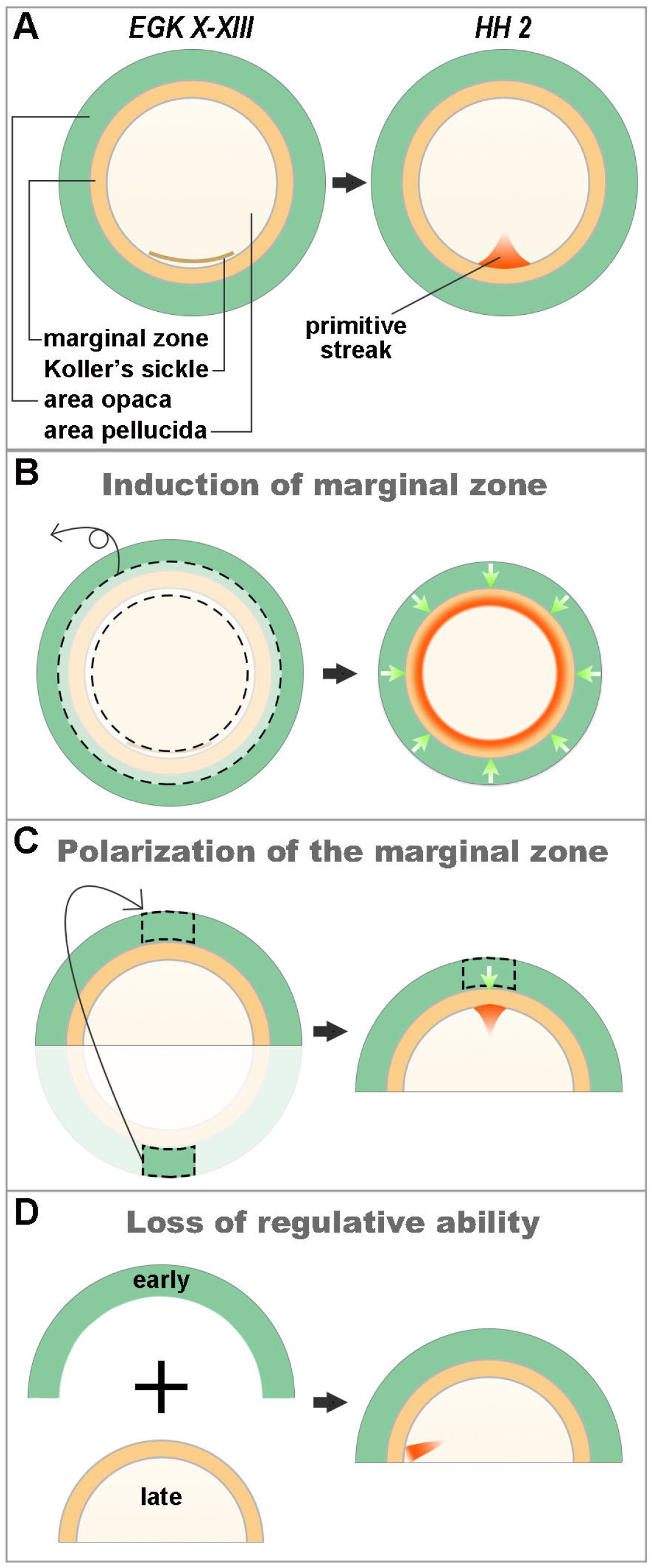
Summary diagram showing the roles of the area opaca on embryo polarity. (A) Anatomy of the early chick embryo at pre-primitive streak (EGK X-XIII) and primitive streak (HH 2) stages. Anterior at the top. (B) The area opaca can induce marginal zone when grafted to the area pellucida (ablation of the marginal zone). (C) A piece of posterior area opaca can induce a primitive streak when grafted onto an isolated anterior half-embryo. (D) Grafting the anterior area opaca of an early embryo (EGK X-XIII) to the inner area (marginal zone plus area pellucida) of a later embryo (HH 2) can rescue the regulative ability of the latter which would normally have been lost at this stage.

Based on a number of classical studies it had been generally thought that the peripheral region of the embryo, the area opaca, plays a role in providing nutrition to the embryo and in maintaining its tension, but that it does not have an instructive role in regulating cell fate or embryonic polarity (Bellairs et al., 1967, Downie, 1976, Khaner et al., 1985, New, 1959, Spratt and Haas, 1960). Contrary to this conclusion, the present study uncovers three separable functions of the area opaca: induction of marginal zone properties, an influence on polarity of the marginal zone, and determining the end of the period during which embryo fragments can “regulate” (re-polarise themselves and form a primitive streak from an isolated fragment lacking the primitive streak forming region) (Fig. 5). We shall discuss these three properties in turn.

### Induction of marginal zone from the area pellucida

When placed directly adjacent to the area pellucida (ablation of the marginal zone; Figs. 1 and 5B), the area opaca can induce marginal zone properties in the latter. Interestingly, this induced marginal zone appears to be entirely posterior in character, all around the circumference, even if the grafted area opaca did not include its posterior part. This raises the possibility that posterior character may represent a “default” condition of the marginal zone, and that this needs to be suppressed by other signals in the normal embryo to restrict posterior character to the appropriate position. This was quite unexpected, as the strong inducing functions of the posterior marginal zone and its expression of cVG1/GDF3, controlled by the transcription factor PITX2 (Torlopp et al., 2014), appeared to reflect an active cell interaction defining this region.

### Polarization of the marginal zone

Ablation of a small piece of lateral area opaca from the edge of an isolated anterior half-embryo biases primitive streak formation to the opposite side (Fig. 2). Additionally, grafting of a piece of posterior area opaca to the anterior region of an isolated anterior half-embryo can induce the formation of a primitive streak (Figs. 3 and 5C). As the primitive streak formation in those experiments is preceded by *cVG1* expression in the marginal zone adjacent to the graft, the area opaca seems to influence the embryo polarity indirectly through the marginal zone, rather than directly affecting the area pellucida. Also, given that the area opaca can polarize the embryo only in the absence of the posterior marginal zone, the polarizing influence of the area opaca is rather supportive and much weaker than that of the marginal zone. This raises the possibility that, before polarization of the embryo (at intrauterine stages) in normal development, the area opaca may have an early role in specifying the marginal zone, which is then polarized. Unfortunately, embryos at these stages are difficult to obtain, to manipulate and to culture.

### Loss of regulative ability

Once the primitive streak forms in the embryo, isolated anterior fragments can no longer regulate: they no longer generate a primitive streak. One possible cause might be loss of competence in the anterior marginal zone in response to the area opaca. However, the anterior marginal zone of HH stage 3 embryos still can respond to cVG1 by generating a primitive streak after grafting a cVG1-expressing cell pellet (Bertocchini et al., 2004), indicating the anterior marginal zone is competent even after primitive streak formation. Here we show that the loss of regulative ability can be rescued by grafting anterior area opaca from a younger, pre-primitive streak stage donor embryo adjacent to the marginal zone of the anterior half of an older, primitive streak stage, embryo (Fig. 5D). This suggests that changes in the signalling properties of the area opaca at later stages are a cause of the disappearance of the regulative ability of the embryo. Formally, these changes could be the gain of an inhibitory signal at later stages; however, the converse experiment of grafting the anterior area opaca from an older (HH2-3) donor to a younger host (anterior half of the area pellucida plus its marginal zone) does not cause a loss of regulative ability in the young anterior half. Therefore taken together, these results suggest that the young area opaca emits positive signals required for the regulative ability of the area pellucida plus marginal zone (and loss of those signals account for the loss of such ability).

### Possible mechanisms: Mechanical tension? Secreted factors (Wnt, BMP?)

What mechanisms might account for the inducing ability of the area opaca and for the loss of support of regulation at later stages? One possibility is that the area opaca may signal to the marginal zone by emitting secreting molecules. WNT and BMP family members are possible candidates. WNT signalling has been shown to collaborate with cVG1 and to be necessary for primitive streak formation (Hume and Dodd, 1993, Skromne and Stern, 2001). One of its ligands, WNT8C, is expressed strongly in the area opaca and is also expressed as a gradient in the marginal zone (Lee et al., 2020, Skromne and Stern, 2001). Once the primitive streak forms at stage HH2-3, *WNT8C* disappears from the area opaca and is only limited to the outmost edge cells (1-2 cell thick) of the area opaca (Lee et al., 2020). Thus, based on its early role on primitive streak formation and its disappearance from the area opaca after primitive streak formation, WNT signalling from the area opaca may be a signal to keep the regulative ability of the embryo. Like *WNT8C, BMP4* is strongly expressed in the area opaca and the marginal zone but with an opposite gradient (strongest anteriorly), and disappears from the area opaca at primitive streak stages (Streit et al., 1998). Although BMP signalling acts as an inhibitor of primitive streak formation and *cVG1* expression, a recent study in the chick embryo suggested that the primitive streak is positioned by the balance between cVG1/NODAL and BMP signals, assessed locally by cells (Lee et al., 2021). Moreover, in mouse embryos, BMP signalling from extra-embryonic tissue induces WNT3 in the epiblast to amplify NODAL signalling and thus position the primitive streak (Ben-Haim et al., 2006).

A second possibility is that tension generated by the expanding edges of the area opaca (the “margin of overgrowth” (Bellairs et al., 1967, Downie, 1976, New, 1959) plays a role. However, embryo fragments lacking the area opaca can regulate (albeit at reduced frequency) ((Spratt and Haas, 1960, Spratt and Haas, 1961) and results in the present paper), and moreover, Spratt’s experiments were performed by growing the embryo in the absence of the vitelline membrane and placed directly (in some cases with its ventral surface downwards) onto a non-adhesive semi-soft agar substrate to which it cannot adhere and which promotes very limited expansion. It is therefore more likely that the influence of the area opaca on the ability of the inner parts of the embryo to regulate is due to secreted signals.

## Materials and Methods

### Embryo harvest and culture and whole mount in situ hybridisation

Fertilised White Leghorn hens’ eggs were obtained from Henry Stewart Farm, UK, and incubated for 2-4 hrs or 13-14 hrs to obtain EGK stage X-XI (Eyal-Giladi and Kochav, 1976) or HH stage 2-3 (Hamburger and Hamilton, 1951) embryos, respectively, at 38°C. Transgenic chick embryos expressing cytoplasmic GFP were supplied by the avian transgenic facility at the Roslin Institute, Edinburgh (McGrew et al., 2008). Embryos were harvested and manipulated (for tissue ablation and grafting experiments described in the text) in Pannett-Compton saline (Pannett and Compton, 1924), then cultured using a modification of the New culture method (New, 1955, Stern and Ireland, 1981) at 38°C for the desired period of time. They were then fixed in 4% paraformaldehyde in phosphate buffered saline (pH7.4) at 4°C overnight prior to whole mount in situ hybridisation as previously described (Stern, 1998, Streit and Stern, 2001). The probes used were: *cVG1* (Shah et al., 1997), *cBRA* (Kispert et al., 1995), and *ASTL* (ChEST817d16, from the chick EST collection, Source Bioscience) (Boardman et al., 2002, Lee et al., 2020). The stained embryos were observed under an Olympus SZH10 stereo-microscope, and imaged with a QImaging Retiga 2000R camera.

### Embryo manipulation

Tissue ablation or excision was conducted in a 35 mm Petri dish filled with Pannett-Compton saline using a fine hypodermic needle or a bent insect pin. Tissue grafting was performed with the embryo on its vitelline membrane after wrapping around a glass ring as described for New culture (New, 1955, Stern and Ireland, 1981). To facilitate adhesion of embryo fragments to each other, they were juxtaposed and any remaining liquid between them was removed by a fine micro-needle pulled from a 50µl capillary, attached to a mouth aspirator. If required, aspiration of liquid was repeated during the first 1-2 hours of subsequent culture. In marginal zone ablation experiments, a thin ring of adjacent area opaca and area pellucida were included with the ablated tissue to ensure that no marginal zone remained. This was confirmed in some embryos using the marginal zone marker ASTL (Fig. 1). To keep track of the orientation of tissues before grafting, its posterior side was marked with carmine powder. After culturing the manipulated embryos for the desired time, the embryo was photographed both before and after *in situ* hybridisation to confirm the orientation of the embryo (as carmine colour is lost during the *in situ* procedure) and to observe morphology more clearly.

